# ChromoTrace: Computational Reconstruction of 3D Chromosome Configurations for Super-Resolution Microscopy

**DOI:** 10.1101/115436

**Authors:** Carl Barton, Sandro Morganella, Øyvind Ødegård-Fougner, Stephanie Alexander, Jonas Ries, Tomas Fitzgerald, Jan Ellenberg, Ewan Birney

## Abstract

The 3D structure of chromatin plays a key role in genome function, including gene expression, DNA replication, chromosome segregation, and DNA repair. Furthermore the location of genomic loci within the nucleus, especially relative to each other and nuclear structures such as the nuclear envelope and nuclear bodies strongly correlates with aspects of function such as gene expression. Therefore, determining the 3D position of the 6 billion DNA base pairs in each of the 23 chromosomes inside the nucleus of a human cell is a central challenge of biology. Recent advances of super-resolution microscopy in principle enable the mapping of specific molecular features with nanometer precision inside cells. Combined with highly specific, sensitive and multiplexed fluorescence labeling of DNA sequences this opens up the possibility of mapping the 3D path of the genome sequence in situ.

Here we develop computational methodologies to reconstruct the sequence configuration of all human chromosomes in the nucleus from a super-resolution image of a set of fluorescent in situ probes hybridized to the genome in a cell. To test our approach, we develop a method for the simulation of DNA in an idealized human nucleus. Our reconstruction method, ChromoTrace, uses suffix trees to assign a known linear ordering of in situ probes on the genome to an unknown set of 3D in-situ probe positions in the nucleus from super-resolved images using the known genomic probe spacing as a set of physical distance constraints between probes. We find that ChromoTrace can assign the 3D positions of the majority of loci with high accuracy and reasonable sensitivity to specific genome sequences. By simulating appropriate spatial resolution, label multiplexing and noise scenarios we assess our algorithms performance. Our study shows that it is feasible to achieve genome-wide reconstruction of the 3D DNA path based on super-resolution microscopy images.

**Author Summary:** The 3D structure of DNA in the nucleus is known to be important for many aspects of DNA function, such as how gene expression is regulated. However, current techniques to localise or determine 3D DNA structure are often indirect. The advent of super-resolution microscopy, at a resolution of 20 *nm* or better can directly visualize fluorescent probes bound to specific DNA in the nucleus. However it is not trivial to associate how many specific stretches of DNA lie relative to each other, making reliable and precise 3D mapping of large stretches of the genome difficult. Here, we propose a method that leverages the fact that we know the sequence of the genome and the resolution of the super-resolution microscope. Our method, ChromoTrace, uses a computer science data structure, suffix trees, that allow one to simultaneous search the entire genome for specific sub-sequences. To show that our method works, we build a simulation scheme for simulating DNA as ensembles of polymer chains in a nucleus and explore the sensitivity of our method to different types of error. ChromoTrace can robustly and accurately reconstruct 3D paths in our simulations.

## Introduction

The primary nucleic acid sequence of the human genome is not sufficient to understand its functions and their regulation. Fitting the 6 billion basepairs or approximately 2 m of double-helical DNA into an approximately 10 *μm* radius nucleus requires tight packing of DNA into chromatin, where about 150 bp of DNA are wrapped around cylindrical nucleosome core particles, which in turn can be tightly packed due to interspersed flexible linker DNA [1]. In addition, each chromosomal DNA molecule occupies a discrete 3D volume inside the nucleus and the arrangement of these chromosome territories is non-random and changes with cell differentiation [2, 3]. This remarkable spatial management of 23 large linear polymer molecules controls crucial functions of the genome, such as gene expression, DNA replication, chromosome segregation, and DNA repair.

Structural biology techniques, such as electron microscopy, crystallography, and NMR have given atomic level insights into the physical structure of the DNA double helix and the nucleosome [4]. In vitro, also higher order structures such as nucleosomes stacked into 11 or 30 *nm* chromatin fibres can be observed and studied at high resolution. However, the existence of regular higher order nucleosome structures in vivo has not been demonstrated under physiological conditions. To date, little direct information is available about the functionally crucial 3D folding and structure of chromatin between the scale of single nucleosomes (approximately 5 *nm*) and the diffraction limit of light (200 *nm*), which can only resolve entire chromosome territories with a size of a few *μm*.

In situ, classically two general types of higher order chromatin organization have been distinguished at a coarser level, euchromatin which tends to be less compact and displays high gene density and activity, and heterochromatin, with a higher degree of compaction and lower gene density and activity [5]. Due to the arrangement of chromatin from individual chromosomes in territories, the majority of DNA-DNA interactions occur in *cis*, while *trans* interactions are more rare and mostly observed on the surface of or on loops outside of territories [6–8]. Within territories and across the whole nucleus euchromatin and heterochromatin are generally spatially separated [9], leading to heterochromatin rich and gene expression poor domains at the nuclear periphery and around nucleoli. Gene expression is intrinsically linked to the 3D structure of chromosomes, chromatin packing densities and the accessibility of DNA by e.g. the transcriptional machinery.

In the last 10 years, biochemical DNA crosslinking technologies based on chromosome conformation capture (3C), have been developed to address the issue of higher order chromatin structure in an indirect manner [10]. These methods have been widely used to measure the average linear proximity of genome sequences to each other in cell populations with good throughput and at kb resolution. The resulting contact frequency maps analyzed with computational models have indirectly inferred principles of genome organization [11]. A major result of these studies, is that chromosomes are organized into domains of 400-800 kb that are topologically associated. These TADs are the smallest structuring units of chromatin above the 150 bp nucleosome level that can be reliably detected biochemically so far. Although good correlations between contact frequency and regulatory elements has been shown for several genes [12], such crosslinking technologies cannot determine the 3D position and physical distances of genomic loci inside the nucleus directly.

Recent developments in light microscopy techniques, collectively called super-resolution microscopy, can determine the position of single fluorescent molecules with a precision of a few nanometers, much below the diffraction barrier. This allows the characterization of previously unobserved details of biological structures and processes [13–16]. First studies have already explored the use of super-resolution microscopy to investigate chromatin structure [17–19], such as the organization of distinct epigenetic states in Drosophila cells [20] that suggested distinct folding mechanisms and packing densities that correlate with gene expression. Dissection of nucleosome organization inside the nucleus in single cells using super-resolution shows that higher nucleosome compaction corresponds to heterochromatin while lower compaction associates with active chromatin regions and RNA polymerase II, and that the spatial distribution, size and compaction of nucleosome correlate to cell pluripotency [18]. While these studies provide first new intriguing insights into chromatin organization they have so far largely focused on single loci without a complete 3D reconstruction of a chromosome or the genome.

However, the resolving power of super-resolution microscopy raises the tantalizing possibility to directly reconstruct the 3D path of large parts of the chromosomal DNA molecule. Super-resolution microscopy can resolve unprecedentedly small volume elements (approximately 20 × 20 × 20 *nm* [21]) inside the total nuclear volume (approximately 8 × 10^−6^*μm*^3^), which will on average contain only up to 2 kb or a few nucleosomes. This fundamental increase in information of the relative positioning of defined loci in the genome can now be leveraged computationally.

This increase in resolution, which enables to distinguish around 60 million volume elements inside a single nucleus, can be combined with any sensitive and site specific fluorescence in situ hybridization (FISH) probe design that allows for spectral and/or temporal multiplexing. Several methods that fulfill these criteria have recently been developed, and fall within two general probe design categories; either a primary imager strand with fluorophore-containing DNA is hybridized to the genome directly [22] or a primary genome-sequence specific DNA probe that facilitates transient binding of the fluorophore-containing secondary imager strand is used (DNA-PAINT) [23]. Our reconstruction algorithm should in principle allow the mapping of the genome sequence in 3D with a resolution of tens of nucleosomes, depending on their local packing density.

Carrying out such large-scale genome mapping studies by systematic super-resolution microscopy will critically depend upon choosing the best design of the necessary chromosome or genome wide fluorescent probe libraries and use sufficient resolution in the employed 3D super-resolution imaging technology. To prove that such studies are feasible and guide their probe design and microscope technology choices, we have developed an algorithm, called ChromoTrace, that uses an efficient combinatorial search to test the theoretical possibility of complete three-dimensional reconstruction of chromosomal scale regions of DNA inside nuclei of single human cells (Fig 1). To thoroughly test our algorithm, we have developed a simulation to model DNA within a geometry similar to that of the human nucleus. Our modeled 3D architecture provides a challenging environment to test our approach, and our ChromoTrace reconstruction algorithm then maps the simulated 3D label positions back to the reference genome. By simulating realistic resolution, label multiplexing and noise scenarios we assessed the algorithm performance for different experimental scenarios. Our results show that ChromoTtace can map the positions of the labeled probes back to the reference genome with very high precision and recall. Importantly, our study shows for the first time that it is feasible to achieve genome-wide reconstruction of the 3D DNA path based on current super-resolution microscopy and DNA labeling technology and defines the required quality of experimental data to achieve a certain bp resolution and reconstruction completeness. This will be invaluable to guide experimental efforts to generate such data sets systematically.

**Fig1.**
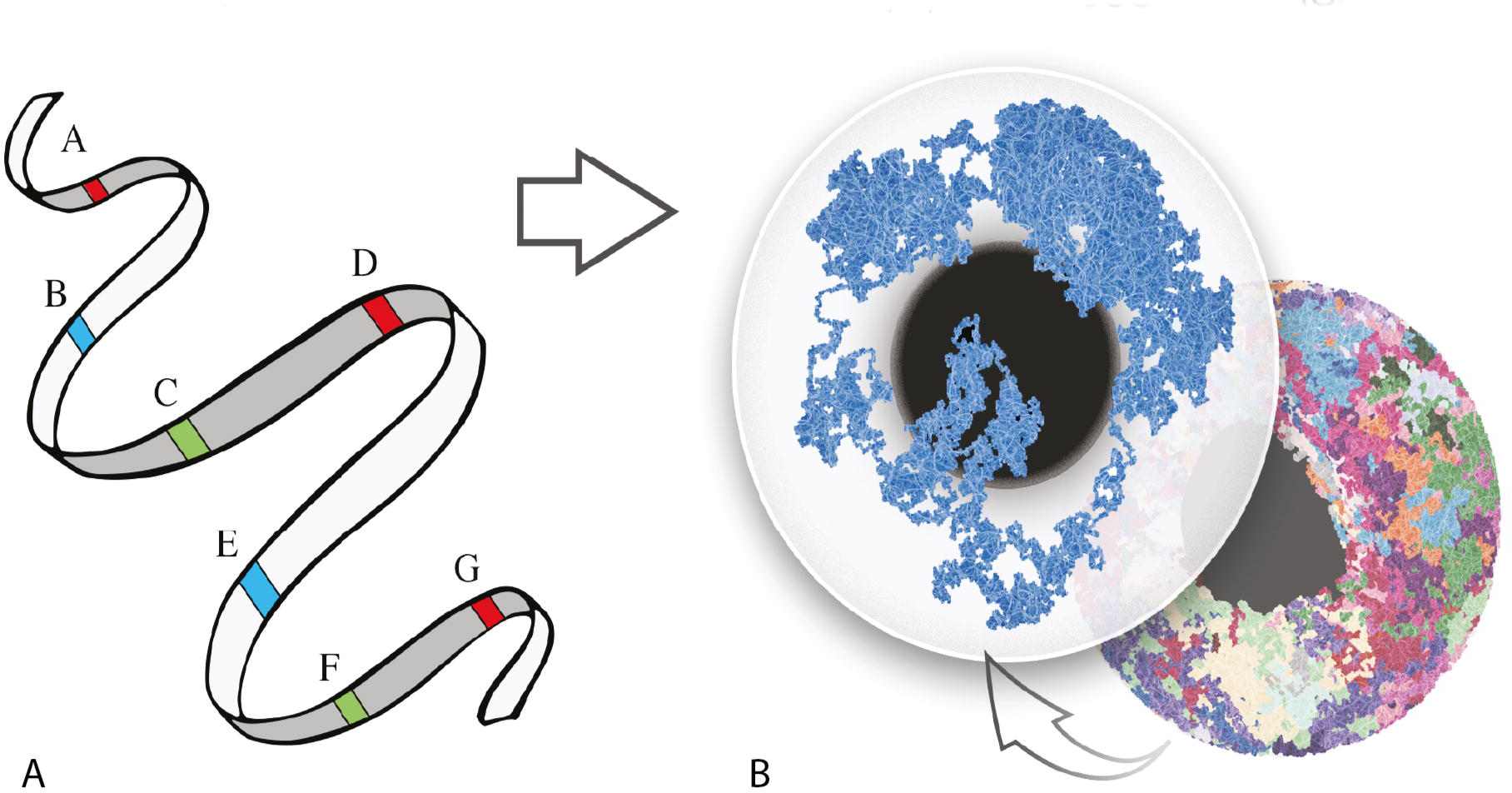
Representation of a chromosome labeling scheme. (A) Linear DNA displayed as a ribbon with six genome regions labelled in three different colors. (B) 3D view of a reconstructed polymer chains. Each globe represents a nucleous and each colored strand within them a polymer chain simulation of a chromosome. The left globe contains only a single chromosome.

## Materials and Methods

### Simulation of DNA in the nucleus

In our simulations we consider each chromosome as a general polymer chain and the whole genome as an ensemble of polymer chains. Each of the polymer chains is modelled as a self avoiding walk (SAW) through a 3D lattice graph. A 3D lattice graph is a three dimensional grid of equally spaced points (from here referred to as nodes), where only the nearest neighbours are connected by an edge and a SAW is a path through a lattice which does not intersect itself. We choose to use SAWs as they are commonly used to model chain-like structures including solvents and polymers, such as DNA [24]. In the following text we use the term *color* equally to either represent different fluorophores, ratiometric labeling with fluorophore mixtures or barcodes, or temporally separated localizations of one or several fluorophores. We generate SAWs through a simple random process. To generate a SAW we pick a random starting node in the lattice that satisfies three conditions.

1. The point is not already part of a SAW.
2. The point is inside the nucleus.
3. The point is outside the nucleolus.

From here the SAW is extended by picking one of the adjacent nodes at random each with equal probability. If the point satisfies the above three conditions then it is added to the SAW; otherwise another adjacent node is picked at random and the conditions checked. This process continues until the SAW reaches the desired length or the SAW becomes stuck and unable to pick any adjacent node. Should the SAW become stuck, we restart this process from another node on the current SAW. Assume the current SAW is of length *i*, we truncate the SAW to length MAX(0,0.8*i*) and begin the process again until the SAW reaches the desired length.

### ChromoTrace Algorithm

In this section we describe ChromoTrace, a new algorithm to identify the 3D structure of chromosomes from a set of labeled points. We begin with an intuitive description of the algorithm and then present the process more formally. The input given to the algorithm is a segmentation file, consisting of a list of (*x*, *y*, *z*) coordinates with associated colors and a labeling file, consisting of a list of genomic locations with associated colors and a distance threshold. The goal of the algorithm is to correctly map the (*x*, *y*, *z*) coordinates to their genomic locations. A brief outline of the ChromoTrace algorithm is given below.

1. Build a suffix tree of the labeling data.
2. Build distance graph of the segmentation data.
  (a) Find all *maximal trivial paths* in the distance graph.
  (b) For each maximal trivial path.
    i. Search the suffix tree and attempt to identify the genomic locations of trivial paths.
    ii. Extend located paths one character at a time until the extension becomes ambiguous.
    iii. Attempt to resolve ambiguous extensions.
    iv. Repeat from Step 2)b)i) until no paths can be extended.
  (c) Remove located paths from suffix tree and distance graph.
  (d) Repeat this process from Step 2)a) until no new maximal trivial paths can be found

### Data Structures

#### Suffix Trees

A suffix tree is a well understood indexing data structure [25, 26] that allows for very fast searching of a subsequence within a sequence. A suffix tree efficiently stores every subsequence of the indexed sequence, for a sequence of length *n* it has exactly *n* leaves and has total size proportional to *n*. The subsequence of the indexed sequence are spelled out as paths from the root of the suffix tree (Fig 2).

**Fig2.**
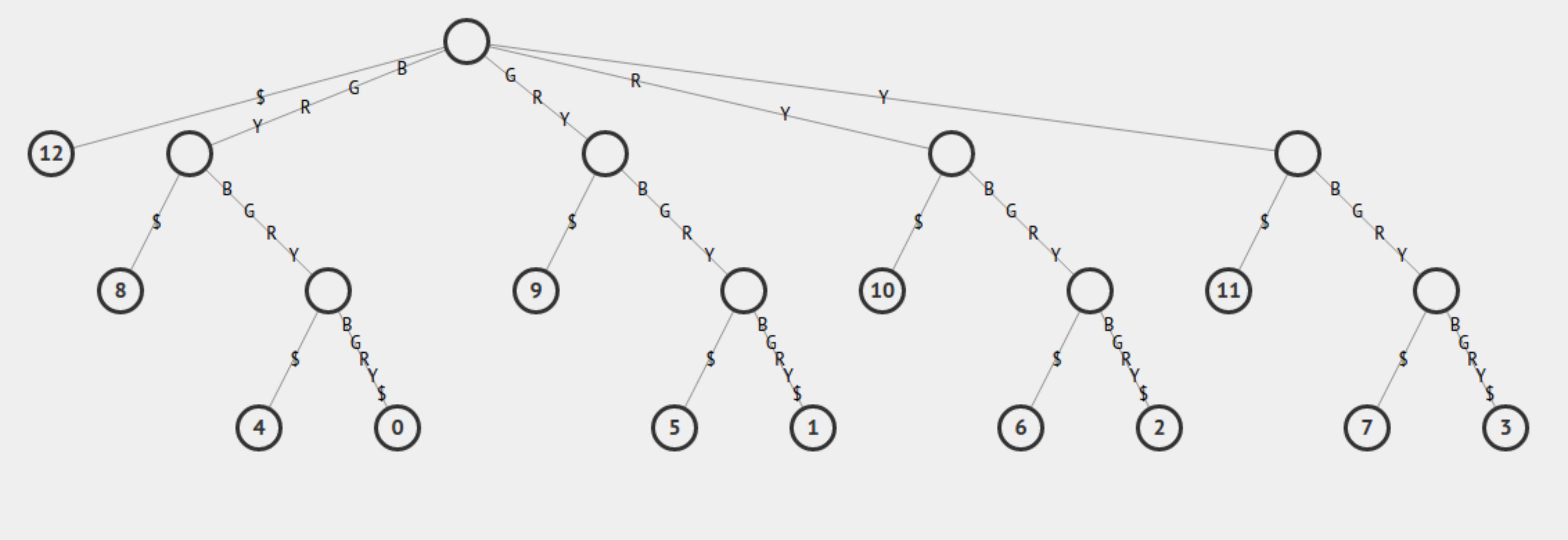
Suffix tree of the sequence *BGRY BGRYBGRY*. Every subsequence of the sequence is spelled out on edges from the root, at the top of the tree, to a leaf node, at the bottom of the tree.

#### Distance Graph

Key to our algorithm is the construction of a graph from the (*x*, *y*, *z*) coordinates in the segmentation file. Each (*x*, *y*, *z*) coordinate will be represented as a node in the graph and two nodes are connected if and only if the Euclidean distance between them is less than a threshold T. T is a user defined value that should be modified depending on spacing of the probe design and resolution of the image. A path in a graph is a sequence of edges, *e*_1_,*e*_2_,…, *e_m_*, connecting a sequence of nodes *v*_1_,*v*_2_,…, *v*_*m*+1_. We define a *trivial* path in the distance graph as a path such that every node connected by the path except for *v*_1_ and *v*_*m*+1_ are required to have exactly two adjacent nodes. A trivial path is maximal if it cannot be extended at either end. More formally we define a trivial path as a path *e*_1_,*e*_2_,…, *e_m_* such that it’s node sequence *v*_1_,*v*_2_,…, *v*_*m*+1_ satisfies the following:

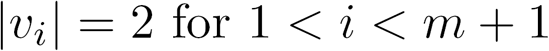

where |*v_i_*| denotes the number of adjacent nodes of *v_i_*. A trivial path is therefore *maximal* if it is also true that for *i* equal to 1 and *m* that |*v_i_*| ≠ 2 or the path forms a cycle.

### Algorithm

After building the suffix tree and processing the segmentation file to build the distance graph we must search the graph to find all of the maximal trivial paths. The set of maximal trivial paths can be found by first storing the number of neighbours each node has and then processing this list. Given the set of maximal trivial paths it is simple to extract the sequence of colors each trivial path represents and to search for this sequence in the suffix tree. If the sequence occurs uniquely in the suffix tree we associate this path with the genomic location found in the suffix tree. Once we have a set of paths mapped to a genomic location we also know which color is expected at the next position in the path. Using this information we explore the distance graph and extend the path with the expected color if there is only one adjacent node with this color. Once we have extended in this way as much as possible there may exist paths where the expected extension is ambiguous. More specifically we may have a path where there are two or more adjacent nodes that are labelled with the expected color. In this situation we find the next L expected colors and search the distance graph for this combination. if there exists an unambiguous extension we add this to the path, otherwise we stop. L is a user defined input where larger values of L will make the algorithm slower but likely increase recall. We repeat this extension process iteratively until no paths can be extended. All of the mapped loci are then removed from the distance graph and suffix tree and the algorithm started again. This entire process is repeated until no more paths can be extended and no more trivial paths found.

## Results

### Simulations of DNA paths in the nucleus

To simulate the results of super-resolved detection of large-scale probe hybridizations, we need to build a model of reasonable packing densities of DNA in the human nucleus. The precise local density of DNA in the human nucleus is surprisingly unclear due to the uncertainty regarding the *in vivo* structure of chromatin. To tackle this problem with reasonable computational complexity at the scale relevant for super-resolution microscopy our simulation uses an intermediate grained self-avoiding-walk (SAW) model for DNA on a grid of points. We use the SAW model to represent DNA as it has been widely used in literature to simulate the structure of polymer chains [24]. The focus of the simulations is to create a challenging environment to test our algorithm and not on creating a realistic simulation of chromatin, although we do use experimentally determined properties where possible.

The nucleus is delimited by the nuclear envelope and contains the nucleolus and DNA. To approximate the structure of the nucleus we used a 3D sphere of 500 *μm*^3^ volume, with an internal sphere of 50 *μm*^3^ volume devoid of polymer chains that represents the nucleolus. In this space we generated 46 polymer chains (two copies for the 22 autosomes and the two sex chromosomes X and Y). Each polymer chain was generated with a length proportional to the chromosome size and the polymer chains were forced to remain inside the simulated nucleus but not allowed to enter the simulated nucleolus. We assume random packing of polymer chains and an average density corresponding to the highest values estimated in human cells, to estimate the sequence reconstruction challenge at a single cell level. Fig 3 shows the packing density and folding characteristics for the ensemble of polymer chains generated by SAWs. Interestingly, although we assumed random packing and no biologically driven heterogeneity in density, the simulation results in a variety of SAW conformations showing broad similarities to known chromatin conformations (i.e., open, fractal and compact). For the remaining text we refer to polymer chains as DNA, refer to each chain by the chromosome it represents in our model and the entire ensemble as a genome.

**Fig3.**
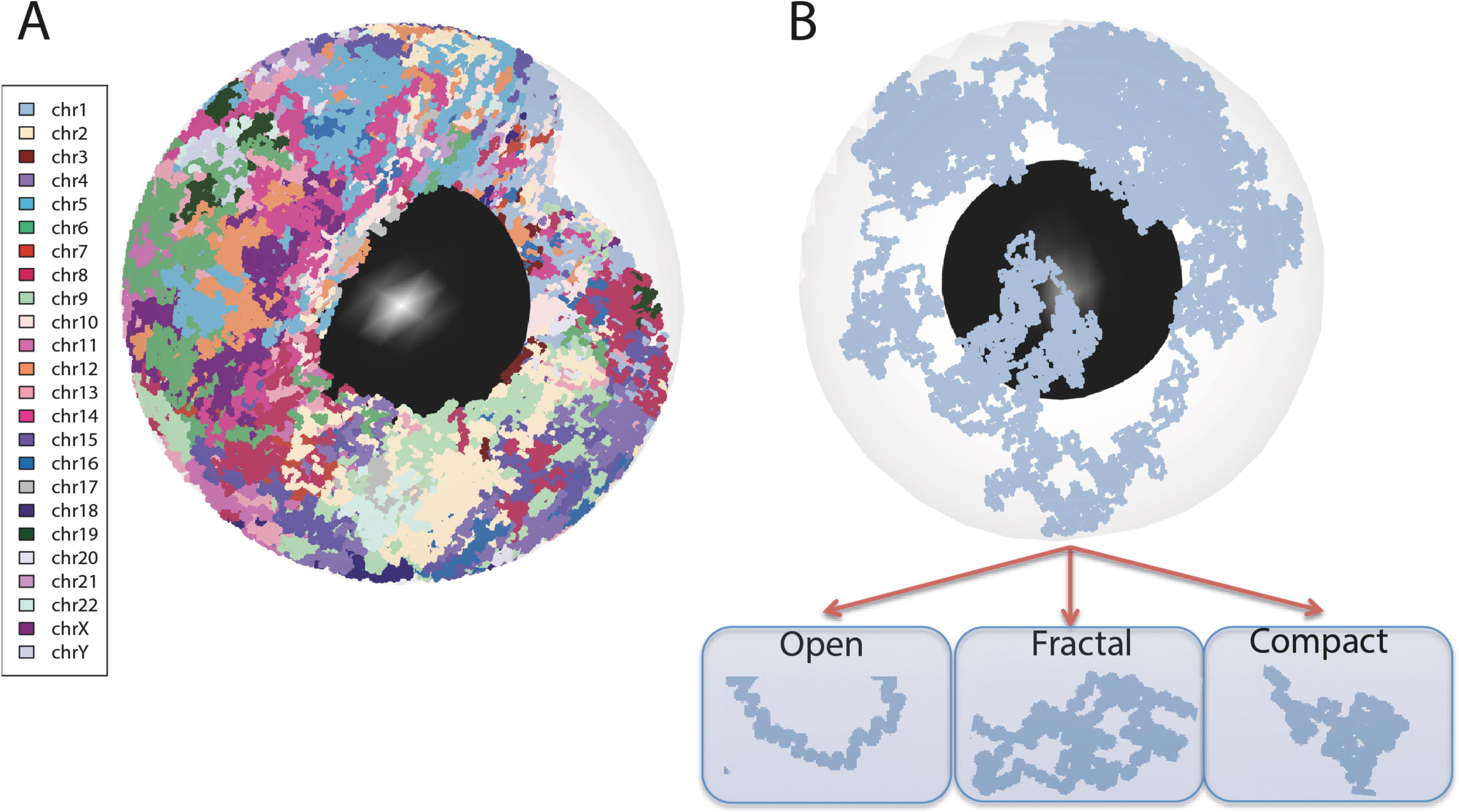
3D view of the simulated genome. (A) An ensemble of polymer chains, each chain, representing two copies for the 22 autosomes and the two sex chromosomes X and Y, is drawn with a different color. Individual polymer chains show a random configuration with a high degree of compactness inside the nucleus. (B) One polymer chain is highlighted showing a variety of polymer conformations with some similarities to, open, fractal and compact chromatin conformations. Each polymer chain is proportional to the size of a chromosome and is labelled as such.

### Testing the mapping of chromosomal DNA sequence to 3D positions of labeled loci

With our ability to simulate DNA in a nucleus, as general polymer chains, we then explored under which experimental conditions, and with what computational methods, the 3D positions of fluorescently labeled genomic loci could be mapped back to the linear chromosomal DNA sequences. Computationally the inputs to the method is a description of the linear labeling of the genome with different colors and the results of the super-resolution image determination, providing a set of the 3D coordinates (*x*, *y*, *z*) and color classification but without the indication of the locus (Fig 4A). The colors where assigned to loci at random, each color having equal probability. The goal of the reconstruction algorithm is to assign each of the in situ loci with a specific (*x*, *y*, *z*) position.

**Fig4.**
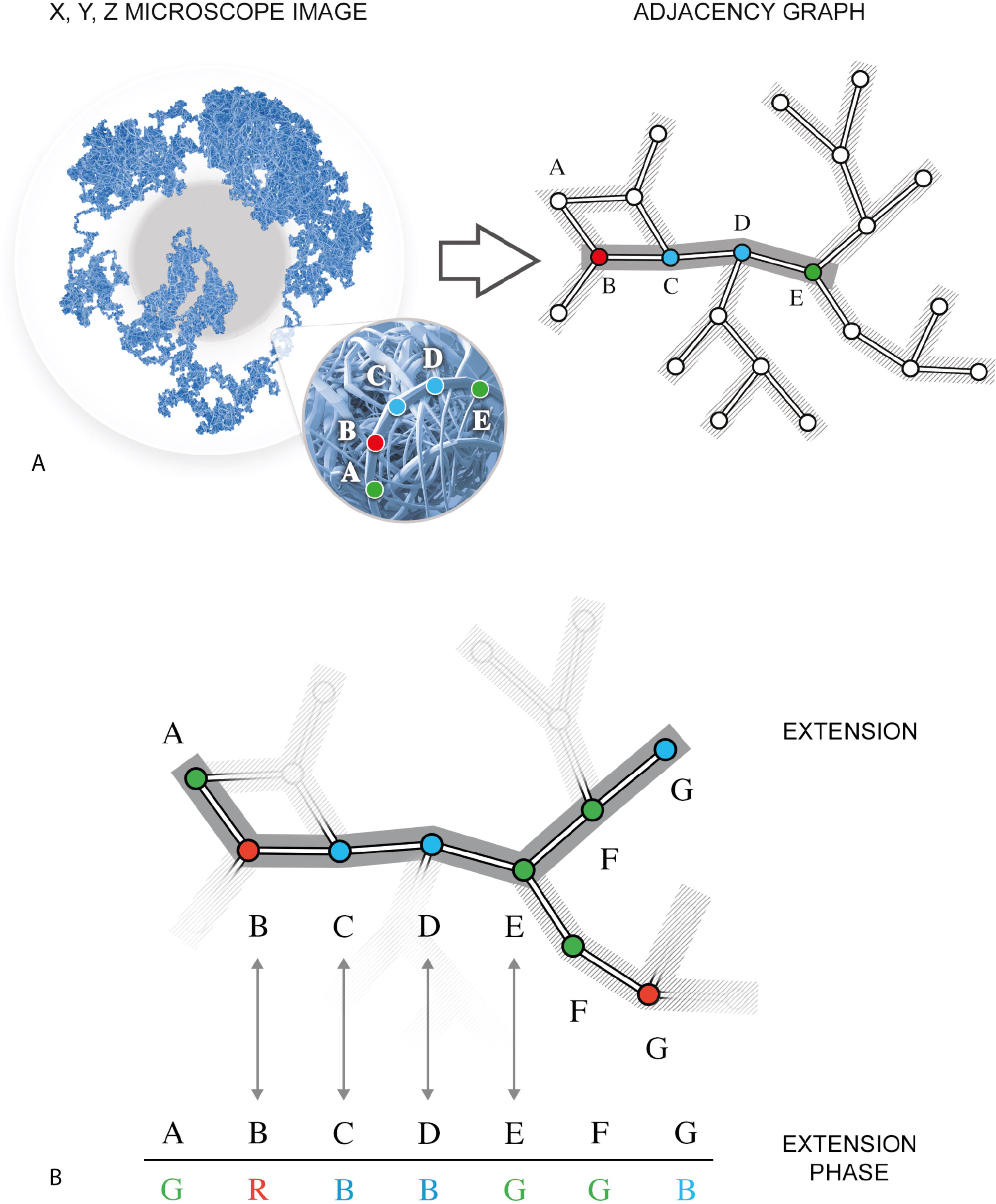
Illustration of the ChromoTrace algorithm. (A) The 3D coordinates that would be obtained from super-resolution microscope imaging are converted into an distance graph. Given the pre-specified linear labeling sequence of green-red-blue-blue-green a trivial path is detected. Note that in the microscopy image the connection between points is unknown and only the colors remain. (B) Diagram of the extension algorithm exploring the ambiguous extension phase.

If we had the same number of colors as loci this task would be trivial, however, the experimental constraints mean that we will have vastly more loci than different colors. Further challenges will occur due to errors in the labeling and imaging experiment. We proposed to solve this problem using the fact that the linear sequence of the probe design dramatically constrains the search space for solutions. Furthermore we can use efficient string based data structures, such as a suffix tree, to efficiently explore compatible places of the design space relative to the 3D space. We named this combined combinatorial exploration followed by expansion the ChromoTrace method (**Methods**).

Our simulations puts us in a position to explore these experimental and technical constraints in a controlled manner, since we can vary the probe design both in terms of number of colors and spacing along the linear genome sequence. Since we know the underlying ground truth of sequence identity and probe color, we can test the hypothesis that the high resolution of 3D position determination and high reliability of color classification provided by super-resolution microscopy should provide enough information to find unique solutions for mapping back probe positions to the linear DNA sequence.

We created probe designs using a regular fixed spacing between probes (in our simulations we use 10.8 kb spacing), resulting in an effective spatial imaging resolution of 4.3 ×10^−5^ *μm*^3^ volume which is well within the limits of super-resolution. We then convert the 3D positions of the simulated imaging data to a graph of potential adjacencies, using a threshold distance of T which relates to the maximum distance between two sequential probe positions in space (10.8 kb) assuming an average compaction of DNA. The resulting distance graph should in theory contain most of the true paths of the probes along the genome plus spurious links of physically close but non-adjacent probes. We then created a suffix tree containing the expected probe colors along the genome, capturing the two possible directions of reading the labels (*p* to *q* and *q* to *p* direction) resulting in a reversible suffix tree with path information for both forward and reverse genome directions. The algorithm then iteratively explores the distance graph to find regions with a unique solution of matching potentially physically adjacent color combinations with the genome sequence (Fig 4A). Once such anchor regions are found, the algorithm has a vastly reduced search space and extends them into the distance graph until it hits regions with high combinatorial complexity (Fig 4B), such as highly compact regions.

To test the performance of ChromoTrace for determining the DNA path through the nucleus we first loaded the labeling file into the reversible suffix tree and jointly searched the suffix tree and distance graph (*x*, *y*, *z*) to find unique sequences of colors found in both. We chose a value for the distance threshold as the value which maximises the number of trivial paths that we find. We performed this analysis for all 100 synthetic nuclear sets, for all 22 probe designs, for all chromosomes separately as well as for the whole genome. Each design is created using a different number of colors ranging from 3 to 24. We choose to use precision (specificity) and recall (or sensitivity) to assess algorithm performance. Recall is the ratio of the number of correctly mapped probes to the total number of probes and precision is the ratio of the number of correctly mapped probes to the total number of mapped probes. Since the ground truth is known *a priori* in our simulation there is no ambiguity in how to measure performance.

Analyzing these 55,000 reconstruction attempts shows that the algorithm is highly precise (mean of 0.99 across all simulations), however the recall rate is much more variable (Fig 5A). This variability can largely be explained by two factors **i)** the number of colors available in the probe design **ii)** the density of probe positions in 3D space. For individual chromosomes the mean recall rate is approximately 0.99 when using a 10 color probe design, however for the same probe design genome wide the mean recall rate drops to 0.64 (Fig 5A). This reflects the increased number of ambiguous sequence paths available when the spatial search space is more densely packed, due to labeled sequences from physically close chromosomes.

**Fig5.**
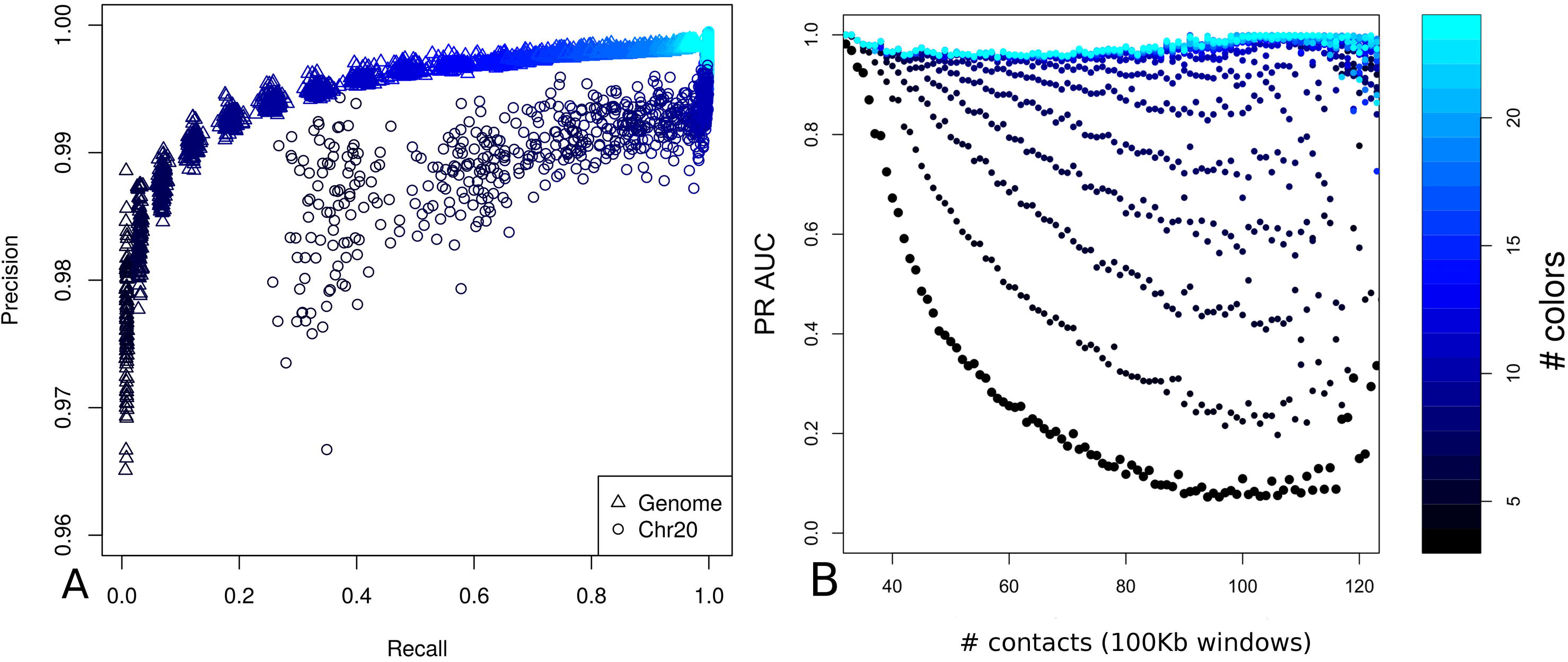
Reconstruction performance for the main simulations. The reconstruction algorithms performance is shown in terms of the relationship between precision and recall given the number of colors in the probe design. (A) Recall against precision genome wide (triangles) and for chromosome 20 (circles). Precision is good for both genome and chromosome scale regions for all the different probe designs whereas recall is much more dependent on the number of available colors and improves as the number of colors is increased. (B) Total number of contacts in 100 kb windows against the area under the precision-recall curve given the number of colors in the probe design.

To assess the reconstruction performance in dependence of the spatial probe position density (i.e. DNA compaction) we show the area under the precision-recall curve values (PR AUC) against the total number of intra-chromosomal contacts in 100 kb windows across all autosomes and for all probe designs (Fig 5B). The contacts are defined as the total number of occupied spaces around each labeled probe, taking into account the grid of points directly surrounding each probe. There is a clear trend for increased PR AUC values for probe designs with a greater number of colors irrespective of DNA density. Across all probe designs there is a marked drop in performance as the DNA density increases, and this drop is much sharper for probe designs with fewer colors (Fig 5B).

For optimum reconstruction it is important to address performance in terms of the completeness of the reconstructed paths. We assessed the length of reconstructed paths in the context of the number of colors available for the labeling design (Fig 6). As expected we see a clear trend towards increased mean and variance of path lengths as the number of colors is increased. Furthermore, the minimum path length across all simulations for each labeling design was 6. The necessity of ChromoTrace to find unique anchors points (trivial paths) before extending further out into the distance graph results in paths never being smaller than 6 points long which is an expected observation when considering the random placement of colors across the simulation space and the global compaction of the chromosomes.

**Fig6.**
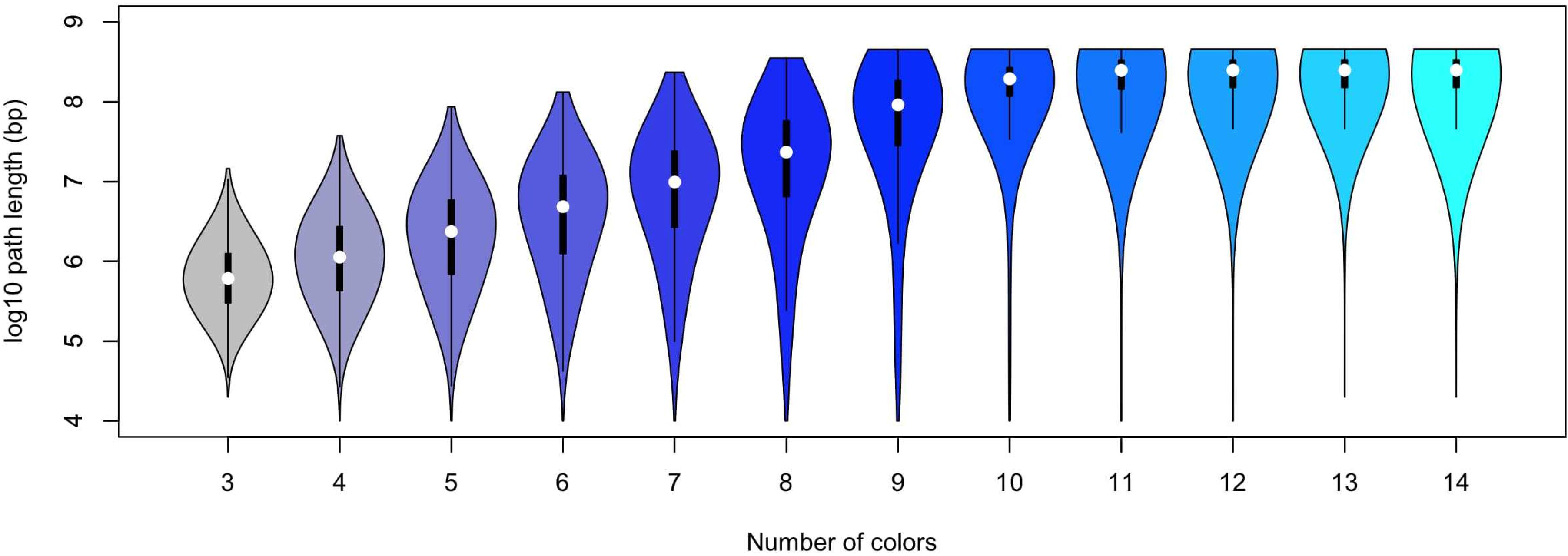
Distribution of path lengths. The relationship between the number of colors and the length of the paths found by ChromoTrace is shown in this plot. A violin plot is shown for each number of colors and the relation to the logarithm (log base 10) of the path length. More colors lead to longer paths and after 10 colors the path length does not increase as recall becomes greater than 0.99. Within the violin plot the first and third quartiles are shown.

### Robustness and error tolerance

Real experimental super-resolution data will contain noise, likely from two major sources, firstly missing probes due to hybridization failure and secondly mislabeled probes, either due to chemical mislabeling or crosstalk between different dyes in the super-resolution microscope. To assess the performance of the reconstruction algorithm in the presence of errors we simulated 99 datasets for each error mode, containing error rates ranging from 1% to 99%, across all 22 probe designs for the 100 simulated nuclear sets, for all chromosomes separately as well as for the whole genome (a total of over 5.4 Million simulations). The errors were added to our simulations at random with the appropriate error rate.

For all probe designs the proportion of mislabeled probes has a dramatic effect on the reconstruction precision and we observe a clear decrease in precision as the proportion of probes with the wrong color is increased (Fig 7A). At 10% mislabeled probes for the 24 color probe designs the mean precision is 0.94 (SD=0.003), dropping to 0.92 (SD=0.006) for 11 colors and to 0.7 (SD=0.003) for 3 colors. Recall rates are even more strongly effected by the proportion of mislabeled probes, starting from a maximum recall rate of approximately 0.99 for the 24 color probe designs with no mislabeled probes, recall rates drop sharply for all probe designs as the proportion of mislabeled probes increases (Fig 7B). At 10% mislabeled probes for the 24-color probe designs the mean recall is 0.85 (SD=0.012), dropping to 0.59 (SD=0.002) for 11 colors and to 0.1 (SD=0.01) for 3 colors. At above 60% of mislabeled probes both precision and recall is too low to be useful. The rapid drop in performance for recall compared to precision is not unexpected considering that ChromoTrace uses exact matching.

**Fig7.**
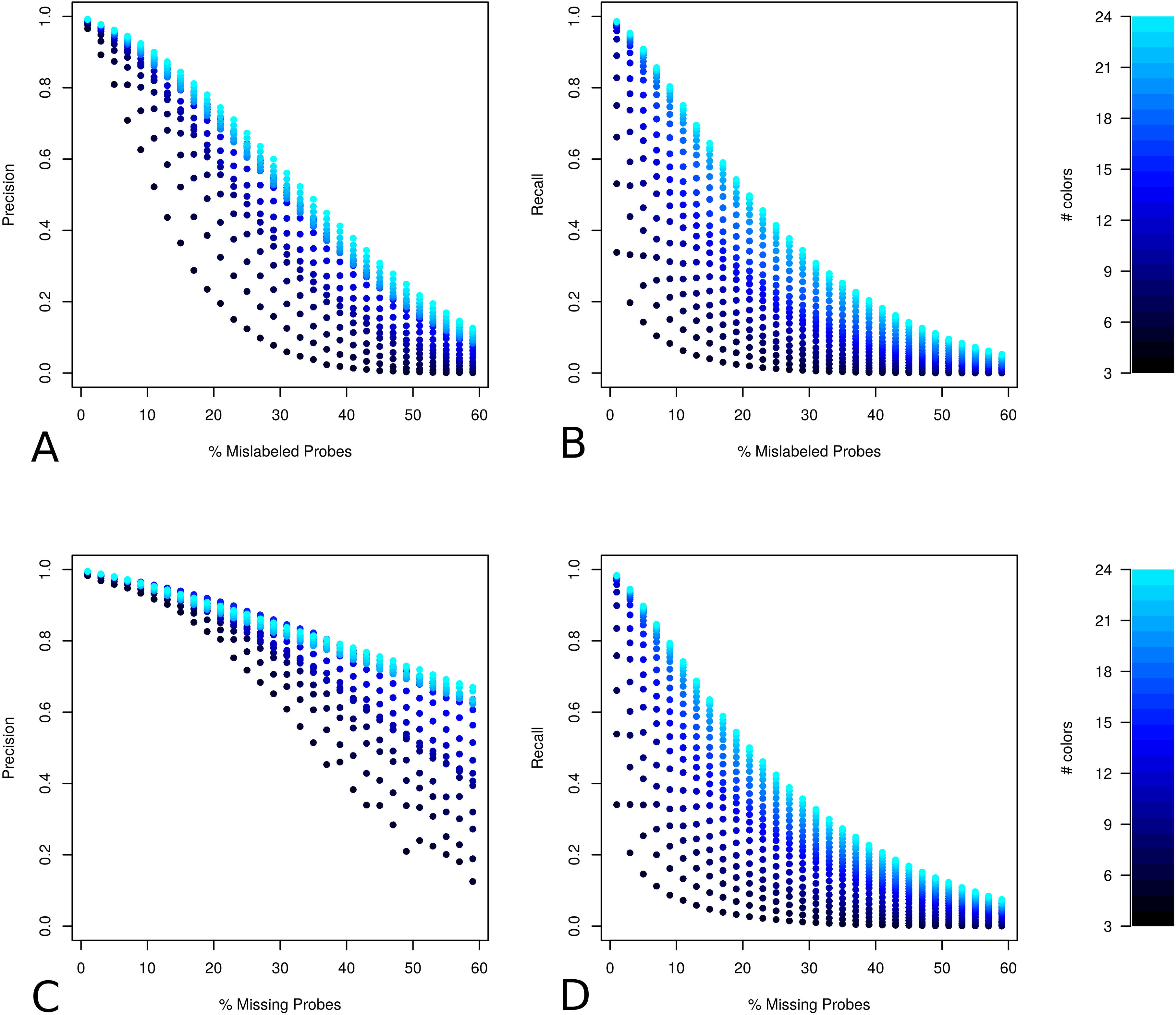
Robustness to missing and mislabeled probes. Relationship between amount of error for two different modes (missing and mislabeled probes) and the overall reconstruction performance given the number of colors in the probe design is displayed in panels A through D. The number of colors in the probe design is indicated using different shades of black-blue. Panels A and C show the proportion of error against precision for mislabeled and missing probe errors respectively and panels B and D show the proportion of error against recall.

For missing probes the relationship between recall and percentage of errors is very similar (Fig 7B and 7D). This is not surprising since either removing or replacing probes with a wrong color in a sequence of colors is likely to stop the extension of correct paths at a similar rate. Precision however, only starts to drop at a much higher percentage of missing, compared to mislabeled probes (Fig 7A and 7C). This suggests that the chance of creating an error in path extension when removing probes is lower than if mislabeled probes are present. If DNA paths were linear in 3D space this would be entirely expected as the distance threshold between sequential probes would ensure that most paths are not incorrectly extended across missing probe locations, while mislabeled probes will not only terminate extension but also cause mismatches to the genome sequence. These results suggest that removing relatively large numbers of probes is unlikely to cause incorrect path extensions across a majority of the simulated DNA space.

Encouragingly even for the probe designs with the lowest numbers of colors (3) precision remains at approximately 0.8 with a missing probe rate of 25%. Furthermore, precision is also relatively robust to mislabeled probe errors, remaining above 0.75 with more than 15% of mislabeled probes for probe designs with greater than 7 colors. As expected recall is far more sensitive to error and there is only a marginal difference observed between the two error modes.

### Differences in DNA packing density

It is unclear how much of the available volume chromatin occupies locally within the nucleus under physiological conditions, but the literature suggests nucleosome concentrations of 140 ± 28 *μM* with nucleosomes every 185 bp in HeLa cells leading to a packing density of 10 % when assuming a nucleosome volume of 1296 *nm*^3^ [27]. To be conservative, our simulation used a higher than average density of DNA, with 34% of the available local volume occupied by DNA (545 thousand points per genome from a 1.59 million point grid). An increased density of the SAWs will result in a harder reconstruction problem, because a higher number of occupied adjacent spaces within the simulation leads to an increase in the number of ambiguous choices for path extension.

To assess the effect of lowering the DNA density we performed additional simulations by omitting the nucleolus and doubling the radius of the nucleus resulting in filling approximately 6.9% of the nuclear volume with DNA (982 thousand points per genome from a 14.1 million point grid). Unsurprisingly these SAWs are less densely packed, an effect that can be visualized by looking at the proportion of adjacent spaces that are occupied, given the distance threshold T, for all labeled genomic locations in the simulations (Fig 8). While in our original simulations the median proportion of occupied spaces around each probe position from the labeling design is 0.52 (Fig 8B), in the lower density simulations this is decreased to 0.37 (Fig 8A).

**Fig8.**
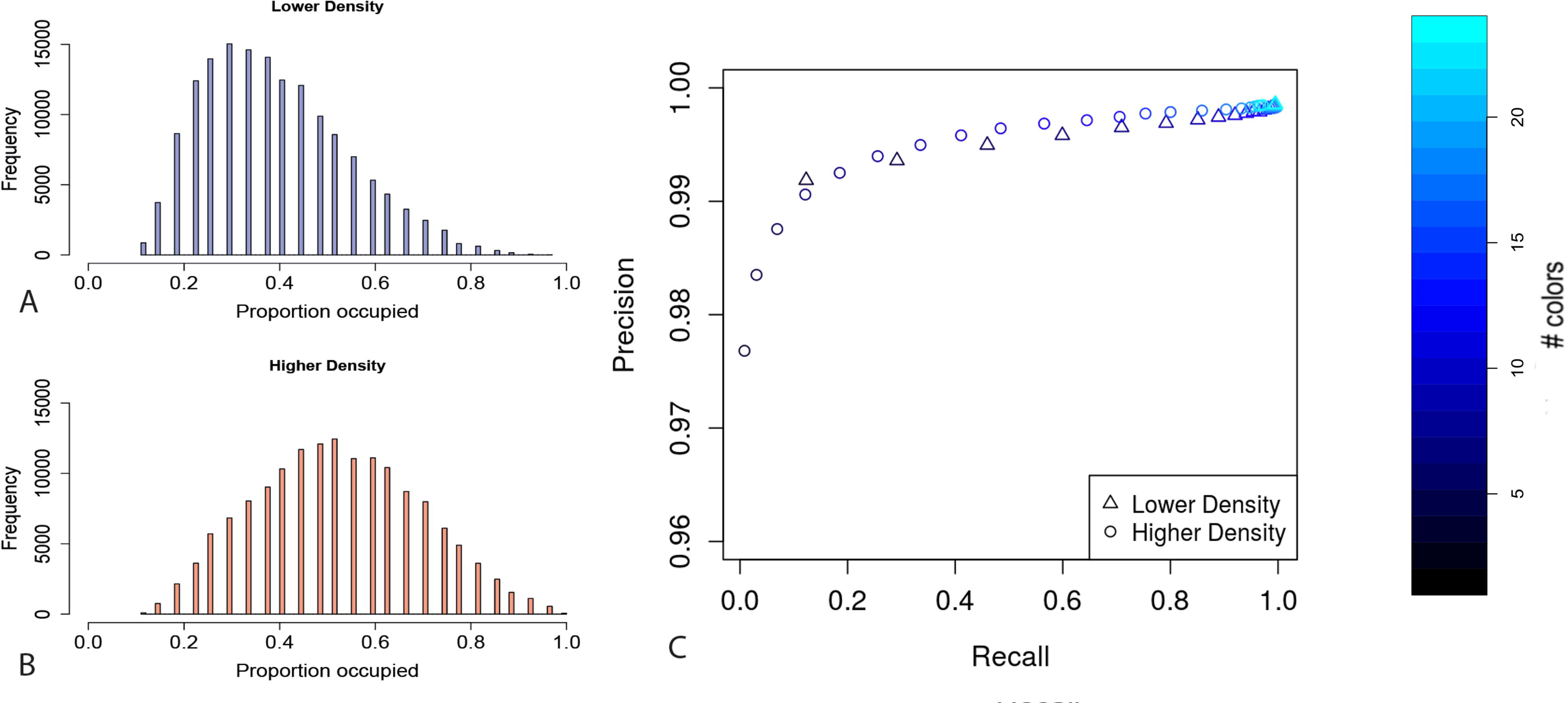
Differences in simulation packing densities. Reconstruction performance when decreasing the packing density of the simulations. (A-B) For all positions across the simulations, the proportion of directly adjacent spaces that are occupied for the new (blue) and original (red) simulations respectively. The distribution is left shifted for the new simulations compared to the original and the median number of occupied spaces is reduced reflecting a decrease in density. (C) Genome wide performance of the reconstruction algorithm for the new (triangles) and original (circles) simulations in terms of precision and recall given the number of colors in the probe design.

To test how this effects the reconstruction performance, we generated 100 synthetic nuclear sets using the approach described above and produced 22 different probe designs containing 3 to 24 colors for the lower density simulations. We then reconstructed using the ChromoTrace algorithm for all synthetic data sets for each chromosome separately and for the whole genome. As expected performance, in terms of both precision and recall, is significantly improved for the less densely packed simulations (Fig 8C). The genome wide mean precision remains high (greater than 0.99) for all probe designs. The difference in recall is much more pronounced with mean recall rates of 0.99, 0.92 and 0.12, compared to 0.97, 0.48 and 0.08, for probe designs with 24, 11 and 3 colors for the lower density compared to the higher density simulations respectively. Importantly when comparing the lower to the higher density simulations the recall rate is improved by a mean factor of 3 across all different color probe designs (Fig 8C). This marked improvement in sensitivity reflects the decreased number of occupied adjacent 3D spaces around each individual probe position and consequently a reduced number of ambiguous sequence path extension choices when lowering the density of the simulated DNA paths (Fig 8A and 8B). Overall across all probes designs the lower density simulations have a genome wide mean recall rate of 0.84 compared to 0.58 for the higher density simulations.

### Simulating Localization Event Profiles

Until this point we have been using simulations containing uniform spacing between adjacently labeled positions (loci), however the distance between adjacent labels in real super resolution experiments will be variable. The main factors effecting this variability are likely to be the lack of absolute uniformity of sequence specific probe spacing along the genome, the super-resolution image localization precision and the probability of effective probe hybridization. To create a more accurate simulation of real experimental data we developed a full simulation of a super resolution experiment. Starting from probe level localization we simulated the results of image acquisition, followed by event clustering leading to observed 3D positions. Importantly this leads to a more varied set of distances between observed loci positions **(Supplementary Information 1)**.

Briefly, starting with the 100 SAWs and the 10 color probe designs, we used each 3D coordinate as the starting point in 3D space and placed 10 probes equally spaced along a single direction (*x*, *y*, or *z*) based on the direction of travel along the walk. The midpoint of each group of probes is the original starting position and each probe was given a 0.3 probability of being missing. Next, for each probe, we simulated a number of localization events (LE’s) drawn from a poisson distribution with a mean of 5 and added error in all directions independently, drawing from normal distributions with standard deviations of 5 *nm*, 5 *nm*, and 15 *nm* for *x*, *y* and *z*, respectively. For clustering these LE profiles we used the DBSCAN (Density-based spatial clustering of application with noise) algorithm [28]. To define the final 3D coordinates for each locus we took the mean coordinate from each direction separately across all LE’s for each cluster that was defined by DBSCAN. These along with their relevant 10 color labeling designs were used as the input to ChromoTrace.

We investigated the result of applying this process to the starting simulations in terms of three different types of error. Firstly, the overall percent of missing loci is approximately 6% for both genomes and chromosomes (Fig 9B), as seen previously the number of missing loci has an extremely adverse effect on the reconstruction performance particularly in terms of recall (see **Robustness and error tolerance**). Next we looked at the percentage of LE’s that were clustered into the wrong locus by DBSCAN, we see that the mean percentage of loci containing erroneous LE’s is approximately 5.8% for both genomes and chromosomes (Fig 9C). The observed position of loci in 3D space whose clusters contain erroneous LE’s are likely to be far less accurate than those whose LE’s are consistent. Moving individual loci around in space is likely to adversely effect the performance of ChromoTrace due to points falling outside of the chosen distance threshold T used in the distance graph. Finally we looked at the percentage of DBSCAN defined clusters that contained LE’s from multiple loci and observe a mean percentage of approximately 1.9% for genomes and chromosomes (Fig 9D). It is reasonable to assume that as the number of unique loci contributing LE’s to a defined DBSCAN cluster increases so the accuracy of the final observed loci coordinates is likely to decrease.

**Fig9.**
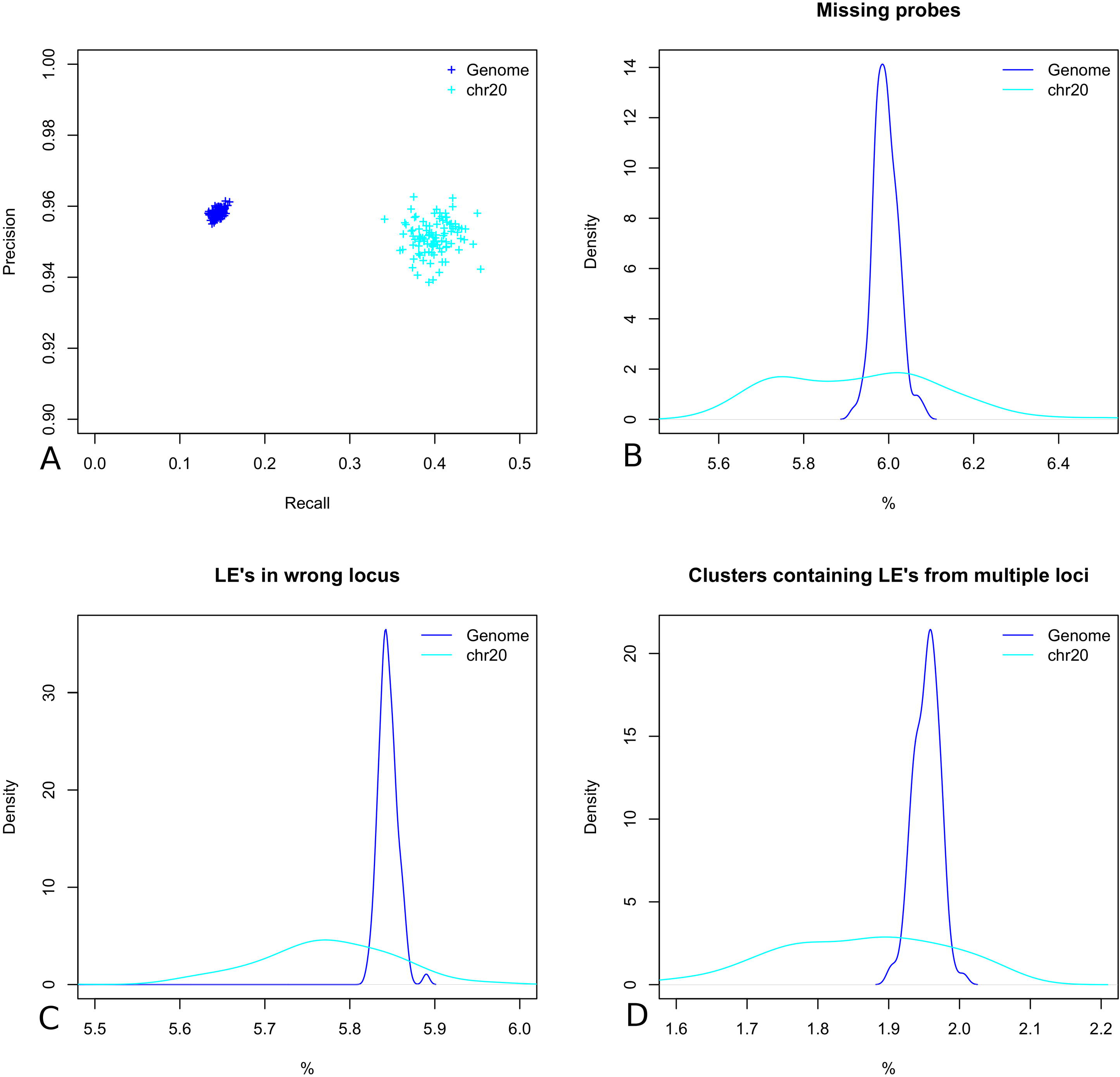
Segmented simulated LE profiles. (A) The reconstruction performance, recall versus precision when running ChromoTrace for whole genome and individual chromosomes. (B) The percent of missing probes across all 100 simulations for all of the polymer chains and a single chain. (C) The percent of LE’s that were clustered into the wrong locus for the whole genome and chromosome 20. (D) the percent of clusters that contained LE’s from multiple starting loci.

We ran ChromoTrace across all 100 simulated super resolution experiments for whole genome reconstruction and for individual chromosomes. Overall the performance in terms of recall was significantly lower than for the original simulations however precision remained high (Fig 9A). The mean precision for both genomes and chromosomes is higher than 0.95 and the mean recall is 0.14 and 0.40 genome wide and for chromosome 20 respectively. The improved recall rates for reconstructing individual chromosomes reflects both the decreased complexity of the distance graph space and importantly a decrease in the number of potential LE’s from different loci that could be incorrectly clustered together by DBSCAN. Here we have chosen challenging parameters for the problem of LE profile loci labeling, profile segmentation and observe a significant decrease in the reconstruction performance achieved by ChromoTrace. However, for chromosome scale genomic regions we are still able to reconstruct approximately 40% of the 3D structure and make very few mistakes with precision remaining above 0.95. The parameters used for simulating these LE profiles are by no means optimal and could certainly be improved when designing real experiments, for example, the number of different color labels could be increased and the rate of missingness improved by using highly specific and locally multiplexed probes. Furthermore the use of DBSCAN in our hands was ‘out of the box’ and we did not attempt to optimise the clustering of individual LE’s from different loci. Improving the clustering of LE’s using DBSCAN or more sophisticated custom algorithms would certainly improve the accuracy of estimated loci coordinates.

## Discussion

We have simulated chromosomal DNA molecules and used a challenging density in the simulated nucleus, however, our simulation is coarse grained and does not at this time take into account a number of known properties of chromatin. One important feature not considered is the known structural heterogeneity of chromatin packing of different genomic sequences, for example eu- and heterochromatic domains or TADs. It is therefore necessary to consider how such structural heterogeneity would affect the reconstruction problem. For a given packing density, such structures should lead to one of two outcomes, firstly that the entire chromosome (or probed region of interest) is overall more compact than simulated, leading to a significantly smaller volume of the chromosome territory. This would effectively reduce the amount of resolvable spatial information present for the reconstruction. Such a result would be disappointing in terms of the reconstruction algorithm, but fascinating in terms of how such chromosomal domains are created and maintained. However, the extended conformation of many chromosomes seen previously [29], along with the distribution of their contacts to the nuclear lamina [28], suggest that overall compaction is an unlikely configuration, except for specific cases such as mitotic chromosomes or the inactive X chromosome. The second outcome is that the more highly packed regions are interspersed with more extended regions. The extended regions would be easier to reconstruct, as the better resolved 3D information will be more accurately able to place these regions to a unique position on the genome. At the extreme of this model one would have a series of resolvable linkers with interspersed globules of packed chromatin that would not be resolvable. In such a scenario integration with the HiC data or other contact maps, whose resolution is good in these more dense regions [30] would be very interesting.

On the other hand, when the density of simulated DNA in the nucleus is lower, the reconstruction improves dramatically in terms of recall. In experimental HiC data if unusual numbers of contacts are observed relative to chromosome size it may be indicative of biological processes effecting chromatin condensation [31]. It is feasible to resolve a large fraction of chromosomal scale regions with a resolution of 10.8 kb and reconstruction at this level would provide very high-resolution chromosomal scale chromatin maps (including the internal structure of TADs, TAD boundaries and inter-TAD regions). Even if the very fine details of high density chromatin structures remain challenging with the currently available imaging technology, the spatial information provided by even partial reconstruction of the chromatin path is certain to increase our understanding of how chromosome folding and partitioning is related to active processes such as gene expression [32, 33].

The other important consideration is the number of distinct fluorescence colors that the reconstruction requires. The number of flourophores compatible with 3D super-resolution microscopy and in-situ hybridization conditions is currently limited to about three dyes that can be reliably spectrally separated if imaged at the same time. Since DNA in situ probes can be coupled to more than one flurophore, combinatorial labeling can create different color ratios. In our simulations, up to 10 colors for simultaneous detection could easily be generated in this manner, however this will also introduce noise due to chemical labeling errors (the chance by which a probe will be labeled with a different color ratio than intended) which would lead to wrong probe assignments. However, since any given color will have only a finite set of possible neighboring mistakes with associated error rates, a substitution matrix of possible errors can be integrated into both the extension phase and exploration phase of the suffix tree [26], changing the formulation of the problem into a likelihood model of seeing the 3D position of probes given a certain path labeling. In addition, recent advances in labeling techniques such as the ‘Exchange-PAINT’ method now allow sequential hybridization and image capture, allowing to separate 10 pseudocolors or more based on a single dye in time [21]. This labeling technology requires long super-resolution image acquisition times, but could massively increase the number of probes available for the reconstruction algorithm. For example, a binary code with 2 colors and 10 labeling rounds could distinguish in the region of 2^10^ labels, which would make reconstruction almost trivial. It is therefore very likely that a well-designed combination of spectral and temporal multiplexing of fluorescent dyes, will make it possible to generate image data with sufficiently large numbers of differently ‘colored’ probes. Therefore it should be possible to optimise data acquisition times with different numbers of colors to allow high resolution reconstruction of the chromatin paths for individual chromosomes within the nucleus. Our comprehensive simulation framework will be valuable in guiding the optimal design of such probes, since it allows to simulate the effect of different designs on the reconstruction performance rapidly in silico.

## Conclusion

In this paper we proposed a novel algorithm, ChromoTrace, to, in theory, leverage super-resolution microscopy of thousands to millions of in situ genome sequence probes to provide accurate physical reconstructions of 3D chromatin structure at the chromosomal scale in single human cells. To test this algorithm we have made simulations of DNA paths in realistic nuclear geometries, and explored different labeling strategies of in situ probes. Our study shows that near complete resolution of a chromosome with 10 kb resolution can be achieved with realistic microscope resolution and fluorescent probe multiplexing parameters. Extensions to this method such as leveraging between nucleus consistency effects and using a likelihood-based scheme will allow even more sophisticated modeling of experimental error sources in the future [34].

There is currently no suitable experimental data to substantiate this work; this is firmly a theoretical exploration of the possibility to achieve this and the constraints any experimental method would need to satisfy for a successful reconstruction. For example, it is clear that minimizing mislabeling is more important than minimizing missing probes. Our simulations are based on known and realistic experimental parameters, where available. We have tested our method under challenging DNA density levels and aggressive error models of missing, misreported data and LE precision. Our algorithm and assumptions are compatible with leading super-resolution techniques; in particular our method assumes isotropic resolution of the probes, which has been shown using methods such as direct stochastical optical reconstruction microscopy combined with interference [21, 35]. Nevertheless real experimental data will likely have properties that we have not anticipated. Some of these properties, such as systematic error behavior, or changes in resolution across the nucleus might hinder our reconstruction. On the other hand, properties such as structured heterogeneity in packing density and cell-to-cell structure conservation are likely to improve our ability to reconstruct. Our reconstructions based on single cell image data are initially most likely to work in a patchwork manner across a chromosome, and will be very complementary to the contact based maps based on HiC or promoter-capture HiC [36]. Combining super resolution imaging and contact mapping should provide fundamentally new insights into chromatin organization and function within the nucleus.

## Supplementary Information

### Supplementary Information 1. Simulation of Localization Profiles

A description of the methods used to simulate localization profiles produced by SRM along with the basic processing used to create the input files for Chromotrace.

